# Gaze tracking of large-billed crows (*Corvus macrorhynchos*) in a motion-capture system

**DOI:** 10.1101/2023.08.10.552747

**Authors:** Akihiro Itahara, Fumihiro Kano

## Abstract

The visually guided behaviors of corvids (Corvidae) are often examined in previous studies because they provide important clues about their perception, attention, and cognition. However, the details regarding how they orient their heads toward the visual targets (or how they use their visual fields when attending to the visual targets) remain largely unclear. This study used a newly established motion capture system to examine the visual field use of large-billed crows (*Corvus macrorhynchos*). Study 1 employed an established ophthalmoscopic reflex technique to identify the visual field configuration, including the binocular width and optic axes, as well as the degree of eye movement. Study 2 used the motion capture system to track the head movements of freely moving crows and examined how they oriented their reconstructed visual fields toward attention-getting objects. When visual targets were moving, the crows used their binocular visual fields, particularly around the projection of the beak-tip. When the visual targets stopped moving, crows frequently used non-binocular visual fields, particularly around the regions where their optic axes were found in Study 1 (close to their visual axes). On such occasions, the crows slightly preferred the right eye. Overall, the visual field use of crows is clearly predictable. Thus, while the untracked eye movements could introduce some level of uncertainty (typically within 15 degrees), we demonstrated the feasibility of inferring a crow’s attentional focus by 3D tracking of their heads. Our system represents a promising initial step towards establishing gaze tracking methods for studying corvid behavior and cognition.

## INTRODUCTION

### What are birds looking at?

Many behavioral studies assume that a focal animal looks at something relevant to its behavioral decisions while gathering critical information for its survival, such as during predator vigilance, prey pursuit, mate choice, individual/species recognition, communication, and social learning. However, inferring an animal’s attentional focus by observing the animal’s head and eye orientations may not be as straightforward as it seems, and this is particularly true for birds (as pointed out by Land, 1999). One reason for this is that, unlike humans and other primates, birds typically do not fixate on a single visual target for prolonged periods but often switch to different eyes and different visual field regions when attending to the same object (Butler et al., 2018; Dawkins, 2002; Kane & Zamani, 2014; Land, 1999). Another reason is that there is a large amount of diversity in birds’ visual field and retinal configurations, such as the size of binocular overlap and the angles of foveal projections, most likely as an adaptation to the species’ unique ecological niche (Fernández-Juricic, 2012; Kulemeyer et al., 2009; Martin, 2007). Thus, although it is a common practice to infer a bird’s attentional focus from its head orientations (eye orientations are typically not tracked except when the bird can wear an eye-tracker as attempted in Yorzinski & Platt, 2014), having detailed knowledge about its visual field and retinal configurations specific to each bird species is crucial for making accurate inferences.

Despite the challenges involved, inferring a bird’s attentional focus should be possible at least with a certain level of accuracy, when there are sufficient clues about the bird and its surroundings. For instance, studies have shown that starlings (*Sturnus vulgaris*) and peafowls (*Pavo cristatus*) orient their foveas, or area *centralis* (functionally and structurally similar to the fovea, except that it lacks the distinctive pit structure characteristic of the fovea), towards a predator image during vigilance (Butler & Fernández-Juricic, 2018; Yorzinski & Platt, 2014). Similarly, terns (*Gelochelidon nilotica*), hawks (*Accipiter gentilis*), and falcons (*Falco rusticolus*, *F. rusticolus/F. cherrug hybrids*, and *F. peregrinus*) orient their foveas towards prey before initiating an attack (Kane et al., 2015; Kane & Zamani, 2014; Land, 1999). Therefore, if we know the species’ visual field configuration, including the angles of foveal projections, and if there is a clearly defined visual target, observing a bird orienting one of its foveas (or any other sensitive spots in its retina) towards the visual target would likely indicate that the bird is looking at it.

However, even when there is a clearly defined visual target, there typically remains ambiguity regarding whether the bird is using its binocular (frontal) field or its non-binocular (lateral) fields. In such cases, additional contextual clues typically help differentiate between these possibilities. These clues include the properties of the object, such as the distance and motion of the object, as well as the behavior of the bird. Specifically, in terms of the object distance, birds predominantly utilize their binocular visual field when controlling their beaks to manipulate objects in close proximity. For instance, pigeons (*Columba livia*) and crows (*Corvus macrorhynchos*) employ their binocular vision when pecking at objects (Goodale, 1983; Matsui & Izawa, 2019), and New Caledonian crows use twig tools (Troscianko et al., 2012). Pigeons and chickens (*Gallus domesticus*) orient their foveas towards salient objects or conspecifics when those visual targets are presented at a distance (Dawkins, 1995, 2002; Kano et al., 2022). In terms of object motion, pigeons visually track slowly-moving objects with their binocular fields and fixate on fast-moving objects with their foveas (Maldonado et al., 1988). In terms of bird behavior, pigeons utilize their upper binocular visual fields during perching flights (Green et al., 1992). Male zebra finches (*Taeniopygia guttata*) direct their binocular fields towards females during courtship displays (Bischof, 1988).

The laterality of eye use also serves as an additional clue. Birds typically exhibit a slight preference for their right eye when attending to detailed features of stimuli and a slight preference for their left eye when attending to more global features of stimuli (see Rogers, 2008 for a review). For example, Australian magpies (*Gymnorhina tibicen*) display a preference for using their left eye to assess a model predator before withdrawing from it, whereas they prefer their right eye when approaching it (Koboroff et al., 2008). European jay (*Garrulus glandarius*), Eurasian jackdaw (*Corvus monedula*), and two species of tits (*Parus palustris* and *P. caeruleus*) prefer to use their left eye when remembering spatial locations and their right eye when remembering object-specific cues in an associative memory task (Clayton & Krebs, 1994). Thus, in these species, the differential use of eyes seems to be generally influenced by both the type of object or task as well as the behavior of the birds, which is considered to be related to their hemispheric specialization (but see Coimbra et al., 2014; Hart et al., 2000 reporting retinal asymmetry of certain species).

In summary, to infer a bird’s attentional focus, the essential information includes the precise orientation of the bird’s head (ideally also its eyes), the species-specific visual configuration, as well as the location of a potential visual target. Additionally, considering additional contextual clues, such as the distance and motion of the visual target, as well as the behavior of the bird, can be informative.

### Gaze of corvids

This study focused on the feasibility of gaze tracking in corvids. Corvids are well-known for their advanced cognitive abilities and complex social behaviors (Baciadonna et al., 2021; Boucherie et al., 2019; Emery & Clayton, 2004; Fitch et al., 2010; Hunt, 1996). Their visually guided behaviors offer valuable insights into their perception, attention, and cognitive processes. For instance, previous studies demonstrated that ravens (*Corvus corax*), like many species of birds, follow the gaze of other individuals (Kehmeier et al., 2011; Loretto et al., 2010; Schloegl et al., 2007). Ravens and several other bird species even follow other individuals’ geometric line of gaze by going behind a visual barrier, suggesting that they take the visual perspective of others (Bugnyar et al., 2004; Zeiträg et al., 2023). Ravens and scrub jays (*Aphelocoma californica*) demonstrate sensitivity to the visual access of other individuals during food-caching events, adjusting their caching behaviors accordingly (Bugnyar et al., 2016; Dally et al., 2006; Emery & Clayton, 2001). However, despite previous studies relying on the birds’ head orientations to infer their attentional focus, the details about how these birds orient their heads toward specific visual targets remain largely unclear. In corvid species, aside from several studies focusing on the visually guided behaviors during object manipulations by crows (e.g., Matsui & Izawa, 2019; Troscianko et al., 2012), no systematic attempt has been made to examine their head and eye movements during object tracking. As a result, there is no technique available to track their gaze directions equivalent to primate eye-tracking (Hopper et al., 2021). This knowledge gap potentially impedes further advancements in the field of corvid behavior and cognition.

### The methods for tracking head movements of birds

There are several techniques available for tracking a bird’s head. The most basic technique involves using fixed standard cameras to record the focal bird’s head angles relative to a visual target in the captured images (Bischof, 1988; Butler & Fernández-Juricic, 2018; Land, 1999; Maldonado et al., 1988). More recent studies have employed head-mounted cameras or eye trackers to investigate birds’ foveal use with the individuals that can tolerate the attachment of such devices (Kane & Zamani, 2014; Tyrrell et al., 2014; Yorzinski et al., 2013). Recently, motion capture systems have been utilized in recent studies to examine the 3D head movements of pigeons by attaching lightweight markers to their heads (Jiménez-Ortega et al., 2009; Theunissen et al., 2016; Theunissen & Troje, 2017).

Most relevant to this current study, Kano et al. (2022) used motion capture systems to track pigeons’ head orientations while they were presented with attention-getting objects. When the object fell within a close distance (roughly within 50cm), they oriented the lower-frontal area of their visual fields to the objects. When the objects were presented at a further distance, they oriented their foveas to the objects. Although eye orientation was not tracked in this previous study, most objects were observed within a few degrees of the foveal projections in the visual field, likely due to limited eye movement in this species, typically within 5° (Wohlschläger et al., 1993).

### The aims of this study

We examined the use of the visual field by large-billed crows (*Corvus macrorhynchos*), a corvid species often studied for their social and visual behaviors (Izawa & Watanabe, 2008; Matsui & Izawa, 2019; Miyazawa et al., 2020). To achieve this, we utilized a newly established motion capture system capable of tracking their head orientations with an accuracy of less than a degree (Itahara & Kano, 2022). Our main aims were to determine which parts of the visual field regions they orient to when presented with an experimentally-defined attention-getting visual target, and what contextual cues influence the differential use of their left and right eyes and distinct visual field regions. By addressing these questions, our larger goal was to establish a gaze-tracking technique in a corvid species. This advancement should prove valuable for future studies on corvid behavior and cognition.

A literature survey indicated that the retinal configuration of this specific crow species is available. Rahman et al. (2006) examined ganglion cell densities in the retina of large-billed crows and determined that the area *centralis* is located near the center of the retina and close to the optic axis in this species. Unfortunately, we did not find any information about the visual field configuration of this species. Therefore, Study 1 examined the visual field configuration of restrained crows using an established ophthalmoscopic reflex technique. Specifically, we assessed the width of the binocular fields and the extent of eye movement in this species.

In Study 2, we tracked the head orientations of freely moving crows using a motion capture system. We then mapped the presented visual targets onto the reconstructed visual fields of the crows. Previous studies have shown that this species of crows relies on their binocular fields during and just before pecking (Matsui & Izawa, 2019). Thus, Study 2 focused on examining how crows use both binocular and non-binocular (lateral) fields when they were presented with objects at a distance beyond pecking range. When using their non-binocular field, it is expected that crows would orient their visual axis (which is close to their optic axis and aligns with the projection of area *centralis*) towards visual targets. However, it remained uncertain how crows differentially use their binocular and non-binocular fields. One possibility is that they use binocular fields only when the object is at a close (pecking) distance, and thus, we may not observe their binocular field use in our experiments. Another possibility is that they use the binocular fields in other contexts as well, such as when the object is in motion (Maldonado et al., 1988). To explore this, we compared crows’ visual field use when the object was in motion or stationary. Additionally, we examined the laterality of eye use in crows. We predicted that our crows might show a right-eye preference because we presented them with small objects of various shapes and colors, which could motivate them to attend to the detailed features of the stimuli.

## MATERIALS AND METHODS

### Study 1 (Visual field configuration)

#### Subjects

Four large-billed crows (*Corvus macrorhynchos*) were used in Study 1 (see Table S1 for the details of the subjects). They were housed socially in a group of 12 crows in an outdoor aviary (130 m^3^) at Uki, Kumamoto, Japan. Nine were captured at a local crow trap (at Gyokuto, Kumamoto, Japan, from January to February 2021) and transferred to our facility courtesy of the farmer, three of which were captured at the study station using a standard crow trap (in February 2021, with permission from the local government, Kumamoto Prefecture; permission no. 02002). All birds were free-floating yearlings at the time of capture. The ages of the birds were estimated at the time of capture based on the pink coloration of their oral cavities (Miyazawa et al., 2020). Study 1 was conducted between September and October 2021. The crows were estimated to be 1–2 years old at the time of the study. The sex of the birds was determined by plucking 2–3 breast feathers and subjecting these samples to standard PCR tests (Fukui et al., 2008). They were provided opportunities to interact with conspecifics and various enrichment objects (dog toys, stones, and branches). They had ad libitum access to water, except during the experiments (approximately one hour on the day of the experiments), and were fed once daily with nutritionally rich foods (dog/cat food, meat, boiled eggs, and wild plants). Three of the four crows (BO_21, BY_21, and YY_21) were used in Study 2. The study protocol was approved by the Institutional Committee of the Wildlife Research Center, Kyoto University (WRC-2021-009A).

#### Ethical note

To capture wild crows, we used one section of our outdoor aviary (64 m^3^) as a crow trap while temporarily opening a one-way entrance (openings with hung wires) on top of the aviary. The local farmer used a standard (commercially available) crow trap (20 m^3^). The crows had ad libitum access to water and fresh meat in the traps. The traps were routinely checked by an experimenter or a local farmer. The crows were kept in the traps for a maximum of one day. None of the crows were severely injured during capture. A few crows had scratches, presumably from the wires at the one-way entrance. We provided veterinary care to the injured crows on such occasions. The crows captured by a local farmer were transported from the capture site to our aviary by car and kept in a dog carrier (or box of the same size) for approximately 1 h. The main experimenter (A.I.) was trained in March 2019 (more than 2 years before the onset of experiments) by another experimenter (F.K.) and a veterinarian to safely catch crows using a net in the outdoor aviary, handle them, and check their health conditions. We caught crows for routine health checks daily and observed signs of distress after they were released back into the aviary, confirming that the catch itself did not seem to incur lasting stress in the crows. The time required to catch the crow was approximately 2 min.

The details of the experimental procedure are described below. We minimized the restraint time for visual field observations (maximally of 1 h daily) and the amount of breast feather sampling (plucking a few breast feathers for sex determination). After the experiment, they were released into the flock for the remainder of the day, and we confirmed that these procedures did not incur lasting signs of distress in the crows. The animals were maintained socially and had free access to food and water except during the experiments. When we found a sick crow, we provided this individual with veterinary care and subsequently isolated it from the flock (but within the visual and auditory access of the flock) in a calm place to rest.

#### Apparatus

We used an established ophthalmoscopic reflex technique (Martin, 1986; Martin & Katzir, 1994). The apparatus comprised a cradle (plastic box stuffed with rubber foam and sponges) and a visual perimeter arm (bent transparent plastic plates; a diameter of 560 mm for observation of the visual field and a diameter of 180 mm for the observation of the optic axes; Fig. 2A,B). The cradle restrained the subject’s body and head by wrapping the subject’s entire body and both legs with soft fabrics/straps and fixing its head with a beak holder (made of metal frames, rubber sheet, and thin wire), padded ear bars, and padded rear-head bar (made of metal bars and silicon). This perimeter arm was a semicircle that extended along the azimuth axis (left and right sides of the bird) and moved along the elevation axis (top and bottom sides of the bird). We positioned the midpoint of the subjects’ eyes at the center of the perimeter arms (the origin of the head-centric polar coordinate system). We set the subject’s lower mandible horizontally in the beak holder (and calibrated the subject’s head in the data analysis such that the line connecting the midpoint of the two eyes and the beak-tip (eye-beak-tip line) matched the crows’ natural perching postures observed in Study 2; see below). The experimenter observed the reflection of the subject’s retina using an ophthalmoscope (LED light; Welch Allyn Ophthalmoscope, Welch Allyn, NY, USA) at each angle on the perimeter arm. The crows were restrained for approximately 1 h. We ensured that breathing was not disturbed when restrained.

#### Procedure

Study 1 was performed over two days for each subject. On the first day, the retinal reflections of the crows were measured while holding the ophthalmoscope on the visual perimeter arm (diameter 560 mm) when the eyes were in a resting position (when no eye movement was induced). To estimate the boundaries of the retina and pecten in each eye, we searched for the edge of the retinal reflection (the angle at which the ophthalmoscope image appeared half-bright and half-dark) in each eye at a step of 1° in azimuth (while smoothly moving the ophthalmoscope along the perimeter arm) and 10° in elevation (while moving the perimeter arm at a step of 10° in elevation). Due to eye movement, we observed the edge of the retinal reflection in several azimuth angles (corresponding to the maximal amplitude of eye movement when the experimenter did not induce eye movement; see Result); we used the maximum angle of azimuth at each elevation as a representative score for each subject, consistent with Troscianko et al. (2012). These procedures were repeated until we completed the measurements of all elevation and azimuth angles, although we skipped a given observation angle when the bird’s body/wings and the apparatus obstructed the view of the ophthalmoscope.

On the second day, we repeated the same procedure while inducing the subject’s eye movement to measure the degree to which the induction of eye movement changed the maximum width of the binocular overlap (or blind area) and to estimate the maximum amplitude of the eye movement from the observed changes. While most previous studies induced bird’s eye movement to estimate the range of the bird’s visual fields when their eyes were either converged (eyes moving to the beak-tip) or diverged (eyes moving to the posterior of the head) (Baumhardt et al., 2014; Martin, 2007), we instead estimated the maximum and minimum ranges of the visual field in any eye direction because of our interest in the 3D eye movements of freely moving crows in Study 2. Thus, we induced eye movement either toward or away from the view of the ophthalmoscope along the perimeter arm (the azimuthal axis; Fig. S1). The measurement interval was changed from 10° to 30° for time efficiency (to maintain a total restraint time of approximately 1 h). Although this compromise in the resolution of measurement along the elevation axis may lower the resolution of the estimated overall shapes of binocular overlap (or blind area) (Potier et al., 2018), we did not change the measurement step of 1° along the azimuth axis. Thus, the interval of 30° along the elevation axis should be sufficiently large to answer our main questions here, namely, the changes in the maximum width of binocular overlap (or blind area) and the maximum amplitude of eye movement estimated from such changes. To induce eye movement in crows, while most previous studies (cited above) used light spots or tapping sounds, we presented a small object (5 × 5 cm) while slightly wiggling it, as we found that this presentation method induced longer eye fixation and made observation easier.

After completing all measurements of the subject’s visual field on the second day, we also measured the angle of the optic axes (when the subject’s eyes were resting). To achieve this, the original visual perimeter was replaced with a smaller perimeter (dia. 180 mm) to make it easier to observe the corneal and lens reflections. We searched for the specific angle at which the reflections from the anterior and posterior surfaces of the cornea and lens (the Purkinje images) were aligned using an ophthalmoscope (two of the four Purkinje images were prominently observed). We performed this at steps of 1° in azimuth and 10° in elevation and repeated this procedure at least nine times per eye for each subject. We used the median value of these repeated measurements as the representative score for each participant. All measurements were performed by the same experimenter (A.I.).

#### Data analysis

A standard visual field reconstruction procedure assumes a hypothetical, infinite viewpoint (Martin, 1984). However, as the ophthalmoscope is placed at a relatively close distance (280 mm, the radius of the 560 mm perimeter arm in our study), a viewpoint correction is necessary based on the distance between the nodal points of the two eyes. This distance was calculated using the distance between the surfaces of the two eyes and the relative location of the nodal point in each eye. The former was calculated for each individual using our custom structure-from-motion application (see the head calibration section of Study 2 for details). Although the latter was not available for this species, a consistent value (2/3 of the distance between the surfaces of the two eyes) was used in a previous study across corvid species (Troscianko et al., 2012; confirmed also from personal communication); thus, we used the same value in this study.

Each subject’s head was calibrated by rotating it in the data by aligning the line connecting the midpoint of the two eyes and the beak-tip (eye-beak tip line) to 0° in azimuth and −10° in elevation (using the 3D coordinates of the eyes and beak-tip calculated from our structure-from-motion application; Fig. 2C). A value of −10° was chosen because the vector connecting the midpoint of the two eyes and the beak-tip points to approximately −10° in the natural perching postures in Study 2 (Fig. S3A). The subject’s binocular field was calculated as the overlap of the left and right visual fields, and the blind area was calculated as the absence of either field. At some measurement points, the edge of the retinal reflection could not be observed owing to obstructions by the equipment or the body and wings of the crow. For visualization purposes in Study 1 and reconstruction of the visual field model in Study 2, we manually filled in the gaps in the measurements based on the shape of the head of the crow. This means that although the subject’s body obstructed the visual fields in our measurement, we did not consider these obstructions as blind spots, considering that the body and wing positions can vary in unrestrained crows. Therefore, these manually corrected values were not used as the main results of Study 1.

### Study 2 (Visual field use)

#### Subjects

A total of 11 large-billed crows were used in Study 2 (Table S1). Study 2 consisted of three experiments. Experiment 1 tested four subjects in June–July 2020; Experiment 2 tested four subjects in May–June 2021; and Experiment 3 tested four subjects in August 2021 (one subject was used in Experiments 2 and 3). An additional six crows were not tested because they habitually removed the markers attached to their heads (see below for details). In Experiment 1, crows were captured at the study station using a standard crow trap (in November 2018, with permission from Kumamoto Prefecture; permission no. 2). Crows in Experiments 2 and 3 were captured by a local farmer or at the study station using a standard crow trap in 2021, as described above. Four of the 11 subjects were housed in an outdoor aviary (64 m^3^, expanded to 130 m^3^ in 2020) socially with a group of five in 2020, and the remaining subjects were housed similarly as described in Study 1. They were estimated to be 1–3 years old during the study period. The study protocol was approved by the Institutional Committee of the Wildlife Research Center, Kyoto University (WRC-KS-2020-12A and WRC-2021-012A).

#### Ethical note

Trapping and housing procedures were identical to those used in Study 1. The details of the marker attachment are described below. Most of the crows immediately tolerated the attachment of lightweight markers. However, we excluded the crows that removed the markers.

#### Apparatus

Crows’ head movements were tracked in a motion capture room built inside the aviary (called the “Corvid Tracking Studio”; W:4 m, D:4 m, H:4.6 m, 73.6 m^3^). This system is optimized to track the fine-scale movements of crows, including head movements, while freely moving and interacting with objects or conspecifics. Unlike other labs that have built a motion capture system applicable to the tracking of various animal species (Nagy et al., in prep), we specialized our system for the tracking of crows because crows typically need long habituation periods in a new room (the room was therefore built inside the aviary) and also because the structure of the room was designed to prevent crows from perching in non-trackable areas (wall angle > 90°) and from damaging the infrared cameras (the cameras were embedded inside the walls; Fig. 1A) (Itahara & Kano, 2022). A perch (W:1.4 m, D:1.8 m, H:2 m) was installed at the center of the motion capture room such that crows could rest and interact with objects/conspecifics well above ground level. Twelve infrared cameras (Flex13, Optitrack, Corvallis, OR, USA; 1280 × 1024 pixels) were installed in the motion capture room; six cameras at the height of 2.5 m were aimed at the ground, and six cameras at the height of 4.5 m were aimed at the perch. The motion capture system recorded the 3D coordinates of the infrared reflective markers at 120 Hz (although our data were smoothed to 36 Hz during processing; see below). Three to five 6.4 mm (in diameter) markers (approximately 0.1 g per marker) were attached to the subject’s head with double-sided tape and glue, which weighed in total less than 1 % of the crow’s weight (680–900 g). Motion capture software (Motive, Optitrack) controlled the cameras and recordings. Before each experimental session, the motion capture system was calibrated with a calibration wand (CW-500, Optitrack) and a calibration square (CS-200, Optitrack). A web camera (BRIO, Logicool, Japan; 3840 × 2160 pixels) was attached to the center of the ceiling to monitor the room and the general behavior of the crows.

**Figure 1.**
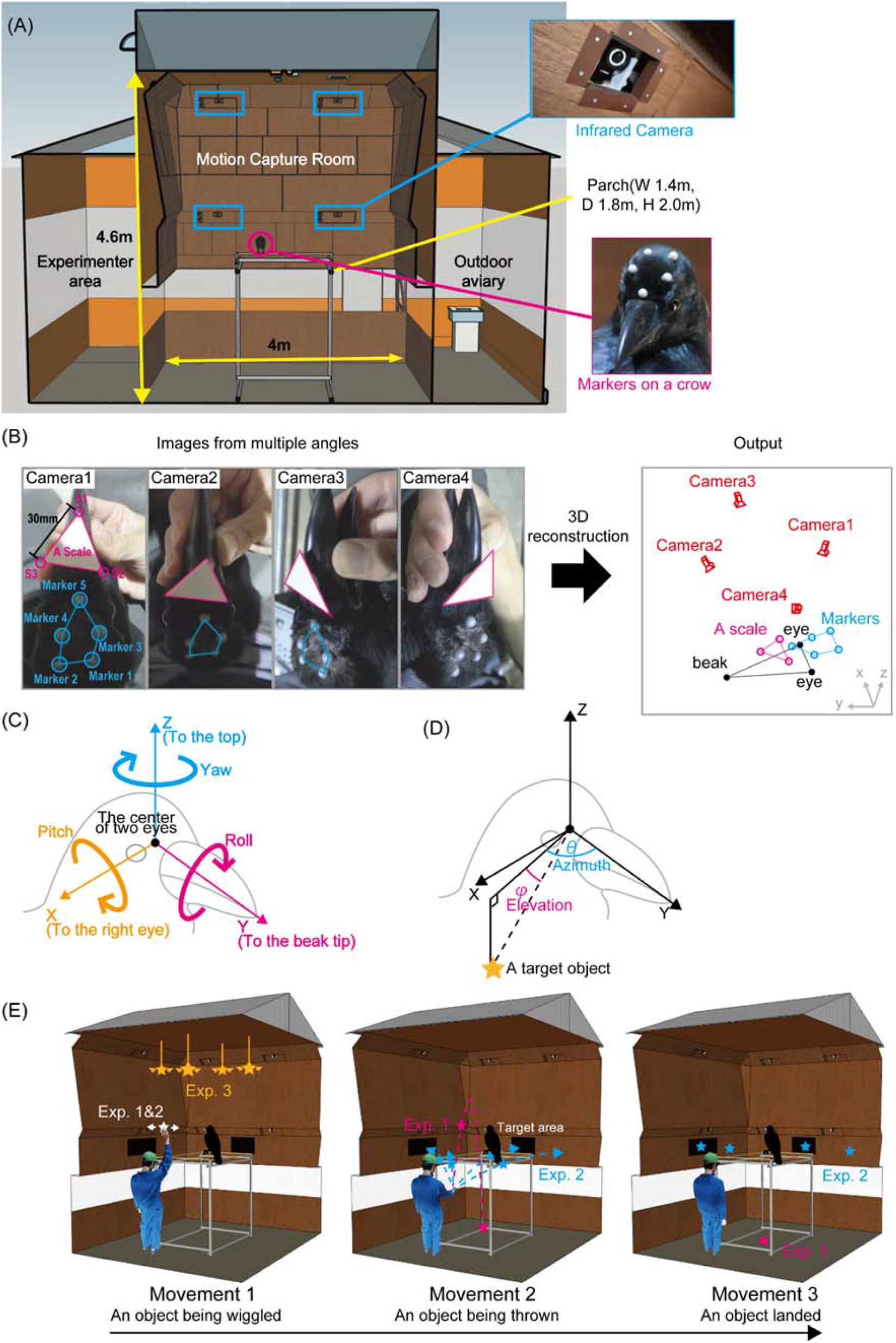
Facilities and procedures of this study. (A) An illustration of the motion capture room (W: 4.0 m, D: 4.0 m, H: 4.6 m). The insets show the infrared camera and the crow’s head with infrared reflective markers. (B) Our structure-from-motion application reconstructs 3D coordinates of key points from 2D images taken from multiple angles. (C) The reconstructed head-centric coordinate system. Thin arrows indicate X, Y, and Z axes. Thick arrows indicate the direction of rotation in yaw, roll, and pitch in this system. (D) The 3D position of a visual target is described in a polar coordinate system (azimuth and elevation). (E) Illustrations of three experiments and distinct movements of visual targets in Study 2. Visual targets were wiggled (Movement 1), thrown to (Movement 2), and landed on (Movement 3) the ground in Experiment 1 and the wall in Experiment 2. Visual targets suspended from the ceiling were wiggled (Movement 1) in Experiment 3.

#### Stimuli and procedure

Before each experimental session (one session per day per subject), an experimenter caught a subject crow with a net in the outdoor aviary, loosely restrained its legs with soft fabrics/strings, placed it on the experimenter’s lap, and attached five 6.4 mm markers to the subject’s head with double-sided tape and glue (these markers were removed after each experimental session to minimize their stress), and calibrated the subject’s head (see below for the detailed procedure). The subject was then released into the motion capture system for approximately 30 min. Only crows that did not attempt to remove the markers participated in the experiments (see the Subject section). Because one crow (GG_20) repeatedly attempted to remove markers when five markers were attached to the head, we reduced the number of attached markers from five to three (the minimum number required to reconstruct 3D movement) for this crow.

In each session, after releasing a subject crow into the motion capture room, we first acclimatized the subject for approximately 2 min until the subject settled on the perch. After acclimatization, in Experiments 1 and 2, the experimenter entered the motion capture room and stood stationarily at a predetermined location for 2 min before presenting the visual targets. As crows are generally sensitive to a human’s direct gaze, the experimenter wore a baseball cap (with reflective markers to track the experimenter’s position) to hide the experimenter’s eyes from the crows. The experimenter wore a wristband with reflective markers on the right hand to track the experimenter’s hand while holding the object. The experimenter then presented various attention-getting visual targets in the subject’s crow. In all experiments, the stimuli were small toys of different shapes and sizes (approximately 2–8 cm in diameter), such as artificial flowers, pocket tissue bags, felt cloths, plastic cups, and kitchen sponges. A 9.5 mm reflective marker was attached to each object to track its position using the motion capture system during the experiments. We performed three experiments in which we varied the presentation height of the stimuli (Fig. 1E).

In Experiment 1, the experimenter removed an object from a waist bag (after holding the hand inside the bag for 10 s), held it above the experimenter’s head, wiggled it for 5 s, and threw it upward; the object followed a parabolic motion and then landed on the ground (approximately at the center of the room) below the eye level of the perched crow. Experiment 2 was identical to Experiment 1, except that the experimenter threw an object onto one of the four target areas on the wall, approximately at the eye level of the perched crow. We attached Velcro tape to the wall and the objects such that the objects could land on the wall. In Experiment 3, four visual targets were suspended from the ceiling (approximately 3 m above ground level) and connected to the experimenter area via transparent threads. The experimenter did not enter the room in Experiment 3 but instead controlled the threads from outside the room (immediately after the 2-min acclimatization of the subject crow). The experimenter wiggled each object individually for 5 s at intervals of approximately 2 min. In all experiments, each session repeated the presentation of a single object (a trial) ten times. Each day, a single session was conducted per participant. In Experiment 1, we performed 20 sessions per bird. Because the results from Experiment 1 indicated a relatively small variation in the crows’ visual responses across sessions, we restricted the number of sessions and performed six sessions each in Experiments 2 and 3. The order of testing for each row was counterbalanced across the days. Each session lasted approximately 30 minutes. After each session, we removed the head markers from the subject’s head (to allow the subject to rest better outside the testing time) and released the subject into the aviary with its social group.

#### Head calibration

The head of the subject crow was calibrated using our custom structure-from-motion application, which reconstructed the positions of the morphological key points (eyes and beak-tips) from the head marker positions. Specifically, before each experimental session, the experimenter loosely restrained the bird (see Stimuli and Procedure) and successively filmed images of the head from four distinct angles using a standard RGB camera (FDR-AX40, SONY, 3840 × 2160 pixels) (Fig. 1B) (Itahara & Kano, 2022). To calibrate this camera (set at a manual focus mode to maintain consistent camera intrinsic parameters), we filmed a printed checkerboard image (64 × 44.8 mm, with each square sized at 6.4×6.4 mm, approximately the size of the crow’s head) from 10–15 distinct angles. During calibration, a triangular scale (30 mm on each side) was temporarily attached to the root of the beak to provide information on the absolute distances between the marked points. The entire calibration process lasted for approximately 30 s.

The RGB camera was calibrated using the filmed checkerboard images in the “Camera Calibrator app” (available in the Image Processing Toolbox) of MATLAB (MathWorks, Natick, USA); this process yielded the camera intrinsic parameters. Using these parameters, the row images were calibrated. In our custom application (GitHub: https://github.com/itaharaakihiro/head_orientation_calibration_app_set), we manually identified the 2D coordinates of the center of the infrared markers, and other key points (the center of the two eyes, the beak-tip, and the three corners of a triangular scale) in each image and then reconstructed the 3D coordinates of these points using the structure-from-motion algorithm.

Finally, from the obtained 3D coordinates of the eye centers and beak-tip, we defined the head-centric coordinate system (comparable to that used in Study 1) with its origin located at the midpoint of the two eyes, the X-axis pointing to the center of the right eye, the Y-axis pointing to the beak-tip, and the Z-axis orthogonal to the X- and Y-axes and pointing to the top of the head. Rotational head movements in yaw, roll, and pitch were defined as the rotation around the Z, Y, and X axes, respectively (Fig. 1C). We then rotated the head-centric coordinate system by −10° in pitch so that the Y axis was oriented to the horizon instead of the beak-tip, as the crows’ eye-beak-tip line (the line connecting the midpoint of the two eyes and the beak-tip) pointed down at approximately −10° in their natural posture (Fig. S3A). We referenced the horizon rather than the beak-tip to reveal the crows’ visual field use with reference to the environment (rather than their morphology). We calculated the angles (azimuth and elevation) at which the visual target was located in the head-centric coordinate system (Fig. 1D).

#### Data analysis

##### Motion capture data processing

In the Motive software, we first defined the rigid bodies (unique arrangements of markers consistent across frames) for the head of the crow and the experimenter’s cap and wristband in a given frame. We then used this information to automatically label all the markers in all frames in the software. Because the software occasionally mislabeled the markers within each rigid body in certain frames (which caused an erroneous orientation of the rigid body), we manually corrected them. Data filtering (noise removal, gap fill, and smoothing) was performed using motion capture software and our custom code (MATLAB), as detailed in Fig. S2. We determined the individual parameters by visually checking the time series of the data to minimize any impossible movement (biologically and physically) and missing values. The customized MATLAB code and sample data are available in our online repository.

##### Saccade filter

A saccade was detected in frames where the axial angle speed exceeded 200°/s (Fig. S5B). This axial angle was calculated from the rotation matrix that converted the head angle recorded in a given timeframe into that recorded in the next frame. Although there was a population of small saccades (shorter than 50 ms and smaller than 5°) in our data, we excluded them from our analyses (and thus included them in the inter saccadic intervals) because they were not distinguishable from the motion capture noise in our data. We then filtered out the moments at which birds made head saccades (8.0 % of all data) because visual processing is likely inhibited during saccades in birds (Brooks & Holden, 1973). Fig. S5C also shows histograms of saccade amplitude, inter-saccadic intervals, and saccade durations.

##### Data extraction

To examine stimulus-driven attention in crows, we extracted the moment crows most likely attended to an object. Specifically, we extracted three critical time windows for object presentation; 1 s (120 frames) following the moment when the experimenter took out the object from the waist bag (defined as Movement 1), 1 s following the moment when the experimenter threw the object (defined as Movement 2), and one second after the moment when the object landed on the ground/wall in Experiment 1 and 2 (defined as Movement 3), and 1 s following the moment when the experimenter wiggled the object suspended from the ceiling in Experiment 3 (defined as Movement 1) (Fig. 1E). At each session, the object was presented 10 times (trials), and we pooled all the data in each session per subject (10 s of data per session). We used a session (day) rather than a single-object presentation (trial) for a subject’s response in the statistical analysis (see below) because the trials yielded too many zeros in our data (which would have unnecessarily complicated our statistical analysis).

##### Visual field definition

We used visual field configurations with the eyes at rest (with no attempt to induce eye movement) in Study 1 (Fig. 2E) to examine the visual field use of the crows in Study 2. We did not consider the projections of the pecten in the analysis of Study 2 as blind spots because their projections could easily be moved by eye movements. To quantify binocular field use, we derived a binocular use score, defined as (binocular looking time – non-binocular anterior looking time) / (binocular looking time + non-binocular anterior looking time). Here, the non-binocular anterior visual field refers to the anterior visual field, between −90° and 90° in azimuth, excluding the binocular areas, as shown in Fig. 5D. To quantify the crows’ use of the left and right non-binocular anterior visual fields, we derived a laterality score defined as (right non-binocular anterior looking time – left non-binocular anterior looking time) / (right non-binocular anterior looking time + left non-binocular anterior looking time). These scores were calculated as the Differential Looking Score (DLS), which is often used in psychology to analyze bias in looking times (Corkum & Moore, 1998; Kano et al., 2019; Senju et al., 2009).

**Figure 2.**
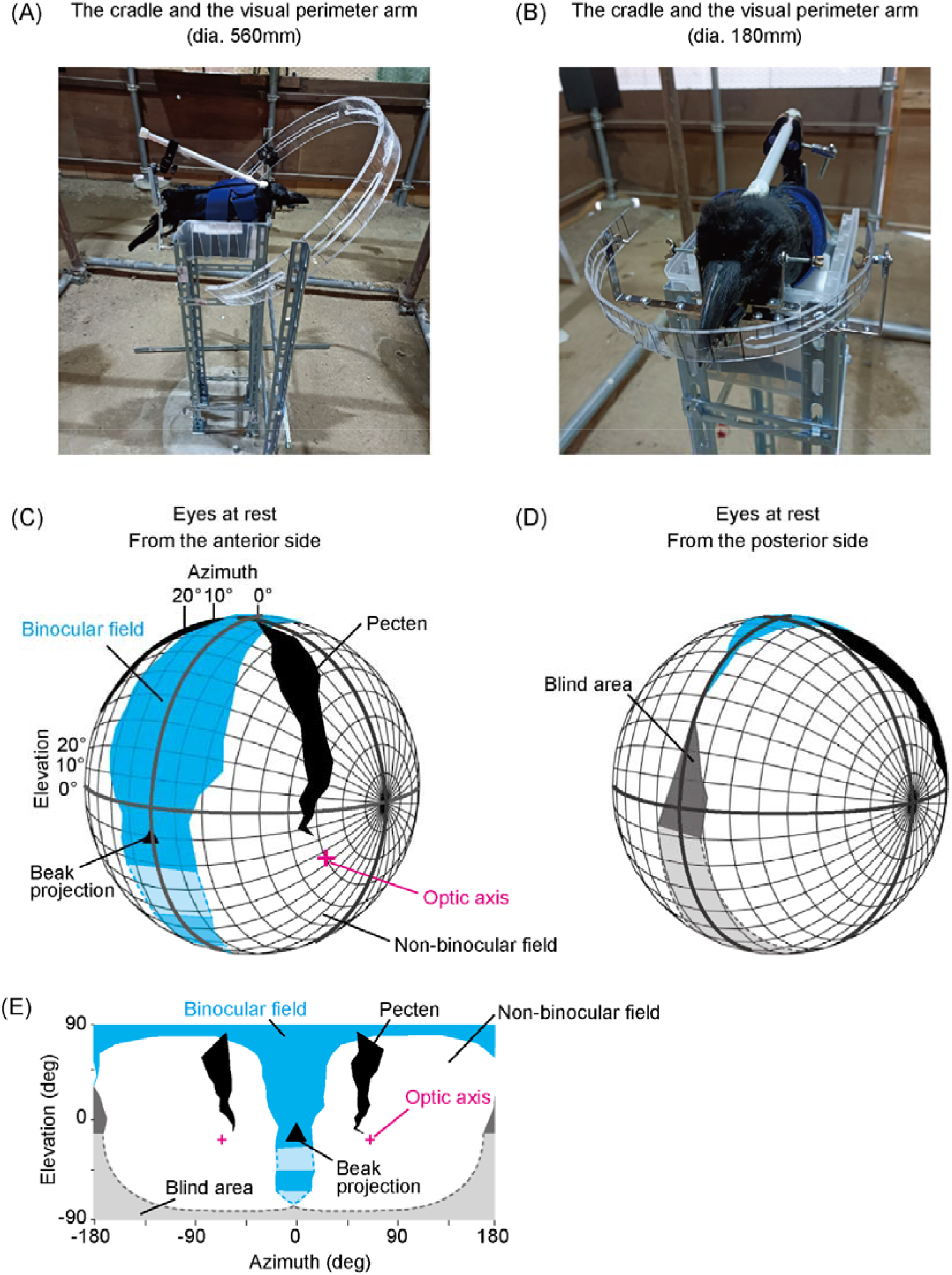
Visual field configurations of large-billed crows. To observe retinal boundaries, the visual perimeter arm of the dia. 560 mm was used (A), and the visual perimeter arm of the dia. 180 mm was used to observe the optic axes (B). 3D illustration of the visual field of large-billed crows in the head-centric polar coordinate system (at 10° intervals) with the eyes at rest (no attempt to induce eye movement). The boundaries of each visual field were calculated as the mean of the four participants. The illustrations show the binocular field (cyan), monocular field (white), blind area (gray), pectens (black), and positions of the optic axes (magenta) viewed from the subject’s anterior (C) and posterior sides (D). The projection of the beak-tip (eye-beak-tip line) is denoted by a filled triangle. Dotted lines and lighter colors denote manually filled gaps in the measurements (when the visual field could not be observed due to obstruction by the equipment or the crow’s body and wing) based on the shape of the head of the crow. The same data were shown to unfold in 2D (E).

##### Statistical analyses

To analyze binocular use and laterality scores, we used R (version 4.1.0) and ran a Generalized Linear Mixed Model (GLMM) with a Gaussian error structure and identity link function using the lmerTest package v. 3.1-3. To analyze the binocular use score, the model included the binocular use score as a response variable, the movement of visual targets as a fixed factor, and subjects and sessions as random effects. Because Experiment 3 had only one type of object movement (wiggling), we compared the results from Experiment 3 with those from Movement 1 (wiggling) of Experiments 1 and 2 (as between-subject comparisons, with the same GLMM structure). To analyze the laterality score, the model included the laterality score as a response variable and participants and sessions as random effects (i.e., with no fixed factor; the significance of the intercept was tested to compare the results with the chance level of zero). In all analyses, we checked the assumptions of normally distributed and homogeneous residuals in the diagnostic plots: histograms of residuals, Q-Q plots of residuals, and residuals plotted against fitted values. It is generally recommended that the levels of random factors be greater than five (Gomes, 2022); however, we had only four subjects (and more than six sessions in all experiments). Although this was recently considered acceptable as long as we were interested in testing fixed effects (Gomes, 2022), we also performed individual-level analyses (the model without the random factor subject) to ensure the observed effects across subjects. To analyze a binocular use score for each individual, we ran the same model omitting the random factor “subject.” We performed a one-sample t-test to analyze each individual’s laterality score. For the group-level analyses, we also checked the model stability for the random factor “session” by removing each session independently and calculating Cook’s distance each time (“influence.ME”) and found that no session was influential (<1) in all analyses. We tested the significance of the model using a likelihood ratio (“drop1” or “anova” in R) comparing the full model with the model without the fixed factor “movement” for the binocular use score or the model without the intercept for the laterality score.

## RESULTS

### Study 1 (Visual field configuration)

The visual fields of the large-billed crows were reconstructed, as shown in Fig. 2C,D. Note that, in our data, the horizon of the reconstructed visual field derives from their natural perching posture (Fig. S3A), and the beak projects into −10° in elevation. When the experimenter did not induce the crows’ eye movement, the maximum width of the binocular field (the maximum range in azimuth at any elevation) was 48.4° ± 1.6° (mean ± SD in four subjects) at an elevation of 16.4° (no SD), and the maximum width of the blind area was 17.1° ± 4.9° at an elevation of 176.3° (no SD). Although we found no SD for the elevation at which the maximum width of the binocular field or blind area was observed (the measurements were performed at a step of 1° in azimuth and 10° in elevation), the calibration of the subjects’ heads (the rotation of the subject’s head based on its eye-beak-tip line) yielded some variations in the calibrated elevation values (SD of ±2.7° for both measurements). The maximum amplitudes of the crows’ eye movement (with no attempt to induce eye movement) were observed as the range of azimuth values in the measurements of the retinal boundaries; 16.1° ± 8.8° for the right eye and 17.3° ± 11.1° for the left eye. The mean angle of optic axes was 62.7° ± 1.5° in azimuth and −32.9° ± 5.0° in elevation for the right eye and 59.4° ± 3.4° in azimuth and −33.8° ± 8.7° in elevation for the left eye (Fig. 2).

Fig. S1 shows the data from when the experimenter induced eye movements of the crows. When the experimenter induced the crows’ eye movement toward the ophthalmoscope, the maximum width of the binocular field was 50.2° ± 4.9° at an elevation of 15.6° (no SD). When the experimenter induced the crows’ eye movement away from the ophthalmoscope, we did not observe the binocular field but instead observed a blind area in front of their head; the maximum width of this blind area was 25.1° ± 11.0° at an elevation of 75.6° (no SD). The maximum width of the blind area behind the head was 24.6° ± 3.8° at an elevation of 165.5° (no SD). As noted above, while we report no SD for the elevation values, the calibration of the subjects’ heads yielded some variations in the calibrated values (SD of ±1.75° for all measurements). The maximum amplitudes of the crows’ eye movement (with attempts of eye movement induction) were 33.9° ± 2.2° at an elevation of 15.5° ± 1.8° for the right eye and 33.8° ± 0.86° at an elevation of 15.6° ± 1.7° for the left eye.

### Study 2 (Visual field use)

An example of the visual field use of a crow is shown in Fig. 3. In this example, the crow moved its head and used different regions of the visual field to track the distinct movements of the visual targets. When the experimenter took out the visual target from the bag and wiggled it (Movement 1), the crow viewed it with its non-binocular anterior visual field. When the object was thrown and followed a parabolic motion (Movement 2), the crow viewed the target with its binocular field and by smooth head movement. When the object landed on the ground (Movement 3), the crow viewed the visual target with its non-binocular anterior visual fields by rotating its head along the roll axis (the Y-axis in our head-coordinate system) with saccadic head movement.

**Figure 3.**
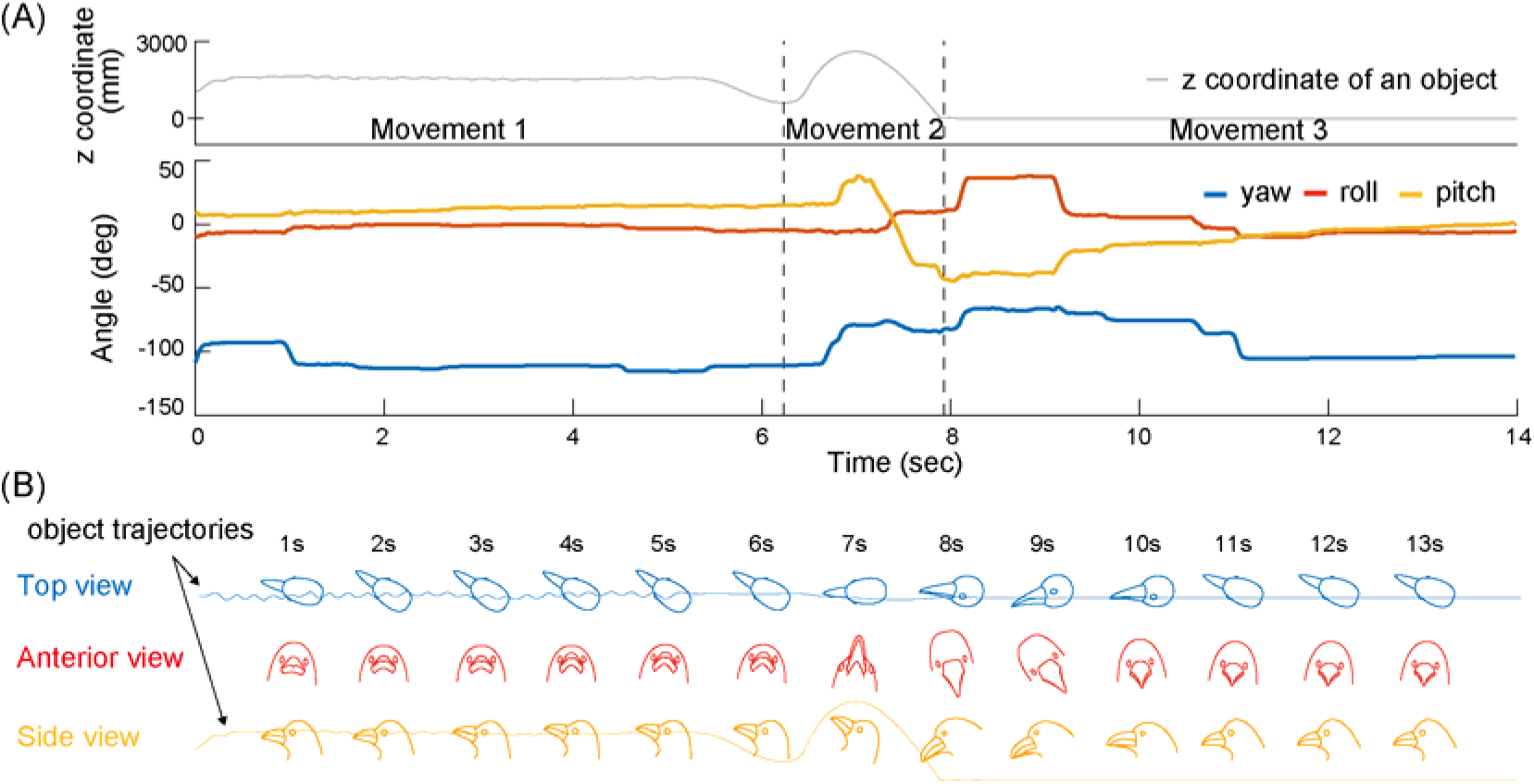
An example of a crow’s head movement when presented with a visual target. (A) The height of an object (in z coordinate) and the rotational movements of a crow’s head (in yaw, roll, and pitch) are shown. (B) The illustration of the crow’s head using the same data.

To visualize the visual field use of all crows, we mapped the distribution of visual targets in their head-centric coordinate system as a heatmap (with the pooled data) for each experiment and target movement (Fig. 4) and mapped the reconstructed visual field from Study 1 (Fig. 2E) onto this heatmap. The heatmaps suggested that the crows mainly oriented their anterior visual field (between −90° and 90° in azimuth) and oriented distinct regions of the anterior visual field to the visual target depending on the target’s movements in Experiments 1 and 2. Specifically, they tended to orient the binocular rather than non-binocular field to the visual target during Movement 1 (wiggled), oriented the binocular field to the visual target even more frequently during Movement 2 (thrown), and then mainly oriented the non-binocular anterior visual field to the visual target during Movement 3 (still). When they oriented the binocular field to the visual target, they tended to orient the regions around the beak projection (the eye-beak-tip line); and when they oriented the non-binocular anterior visual field to the visual target, they tended to orient the regions around where we found the optic axes in Study 1. In Experiment 3 (wiggling the object on the ceiling), the crows tended to orient the binocular field to the visual target. However, the distribution of visual targets seemed more moderate overall than in Experiments 1 and 2, presumably because the visual targets in Experiment 3 attracted the crows’ attention less strongly than those in Experiments 1 and 2.

**Figure 4.**
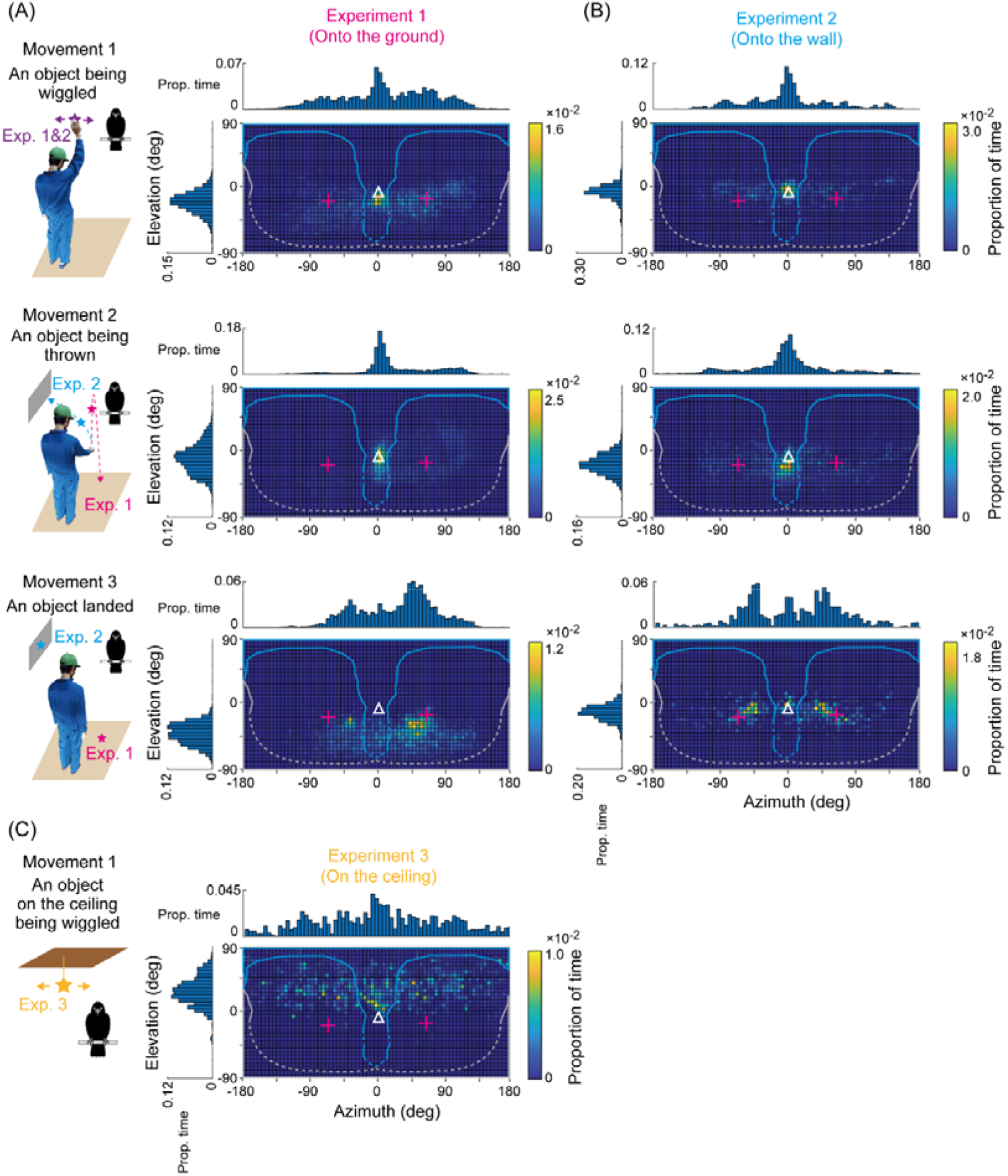
The distribution of visual targets in the crows’ visual fields. Distribution of visual targets in crows’ visual fields in Experiments 1 (A), 2 (B), and 3 (C). The three rows in A and B show the distinct movements of the visual target, with an illustration on the left. Each bin was 5° × 5° and indicated the proportion of time when visual targets were observed in this bin. In addition, histograms (binned at 5°) plot the same data in azimuth and elevation. The reconstructed visual field from Study 1 (Fig. 2E) was superimposed onto each heat map. The plus marks denote the optic axes in Study 1. Blank triangles denote the projection of the beak-tip (eye beak-tip line).

To quantify the crows’ differential use of binocular fields across the targets’ movement and experiments, we derived the binocular use score as (binocular looking time – non-binocular anterior looking time) / (binocular looking time + non-binocular anterior looking time); note that non-binocular anterior region refers to the anterior region (the azimuth of ±90°) excluding the binocular area. We ran a Generalized Linear Mixed Model (GLMM) with this score as a response and the movement of the visual target as a fixed factor. As random effects, we included the random intercept of the subjects and sessions and the random slope of the movement of the visual target (for both subjects and sessions). The binocular use score varied depending on the movement of the visual target in Experiments 1 and 2 (likelihood ratio test, χ^2^ (2) = 8.88, *P* = 0.0118 for Experiment 1; χ^2^ (2) = 13.31, *P* = 0.0013 for Experiment 2); as Fig. 5 shows, it was the largest in Movement 2, followed by Movements 1 and 3. We performed the same analysis (the same GLMM without the subjects as a random effect) at the individual level and found that all crows showed similar results (the maximum *P*-value was 0.0070). As Experiment 3 contained only one movement of the target, we compared the binocular use score of Experiment 3 with that of a similar condition, namely Movement 1 (wiggled) of Experiment 1 and 2, using a between-subject analysis (with the same GLMM structure). No significant differences between the experiments were found in these analyses (likelihood ratio test, χ^2^ (1) = 0.99, *P* = 0.3198 for Experiments 1 and 3 and χ^2^ (1) = 0.37, *P* = 0.5405 for Experiments 2 and 3).

**Figure 5.**
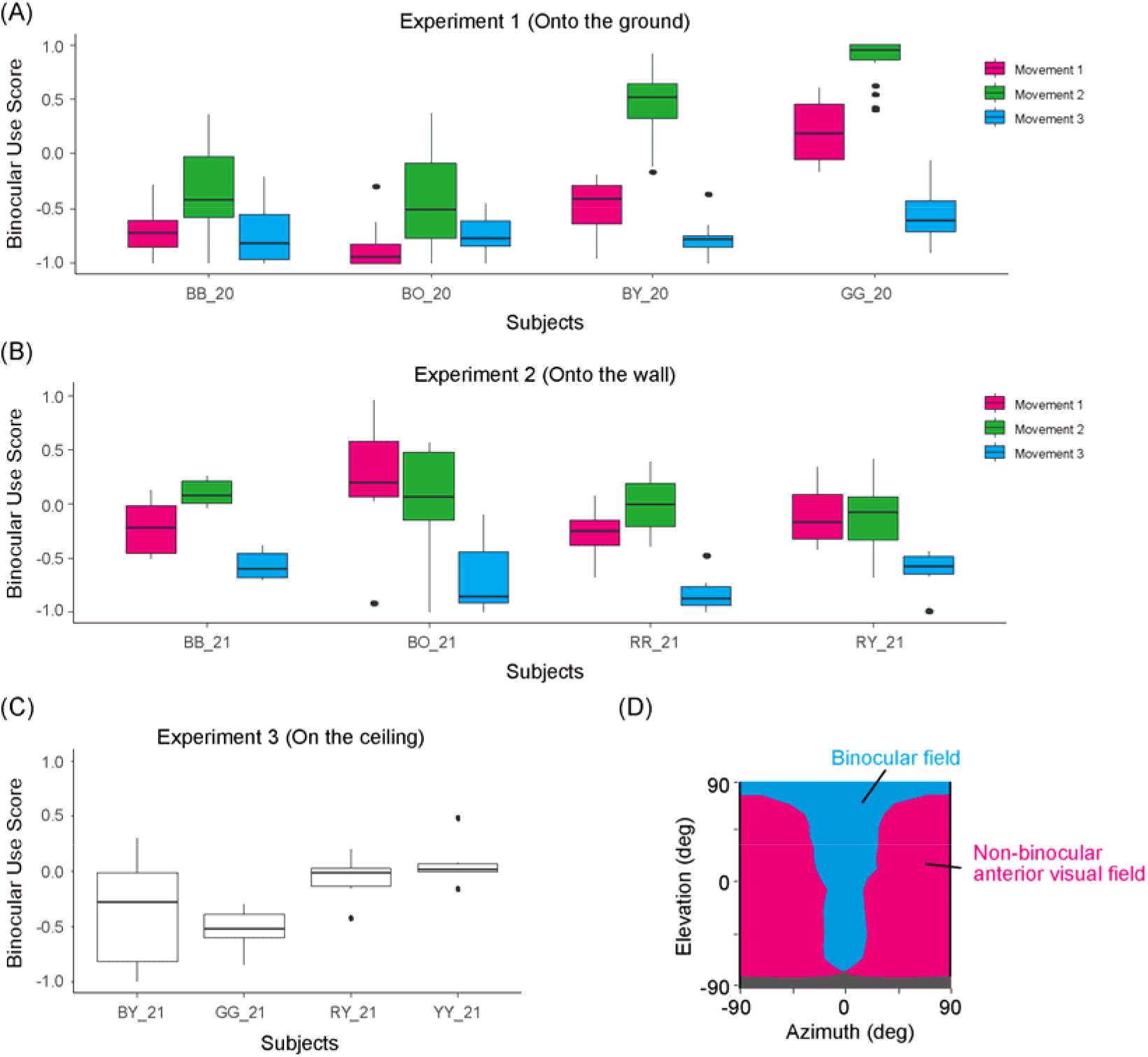
Binocular use scores in each experiment. Binocular use scores were defined as (binocular looking time × non-binocular anterior looking time) / (binocular looking time + non-binocular anterior looking time) in Experiments 1 (A), 2 (B), and 3 (C). Box plots show the median, interquartile range (IQR), and 1.5 × IQR, with outliers plotted individually. In addition, the 2D visual field map created in Study 1 was used to calculate the binocular use score (D); the binocular field, non-binocular anterior visual field, and blind area are indicated in cyan, magenta, and gray, respectively.

We also examined the laterality of eye use by defining the laterality score as (right non-binocular anterior looking time – left non-binocular anterior looking time) / (right non-binocular anterior looking time + left non-binocular anterior looking time). We used data from Movement 3 of Experiments 1 and 2 because the crows mainly used non-binocular anterior visual fields in this condition. We ran a GLMM with this score as a response and the same factor/effect structures as described above. The crows showed a right-eye preference in Experiment 1 (likelihood ratio test, χ^2^ (1) = 7.04, *P* = 0.0080) and a similar non-significant tendency in Experiment 2 (χ^2^ (1) = 3.36, *P* = 0.0667) (Fig. 6). For the former (Experiment 1), we also tested the laterality bias in each individual and found that three out of the four crows showed significant right-eye preferences (The *P*-value for GG_20 was 0.1594, whereas the maximum *P*-value was 0.0094 for the other crows).

**Figure 6.**
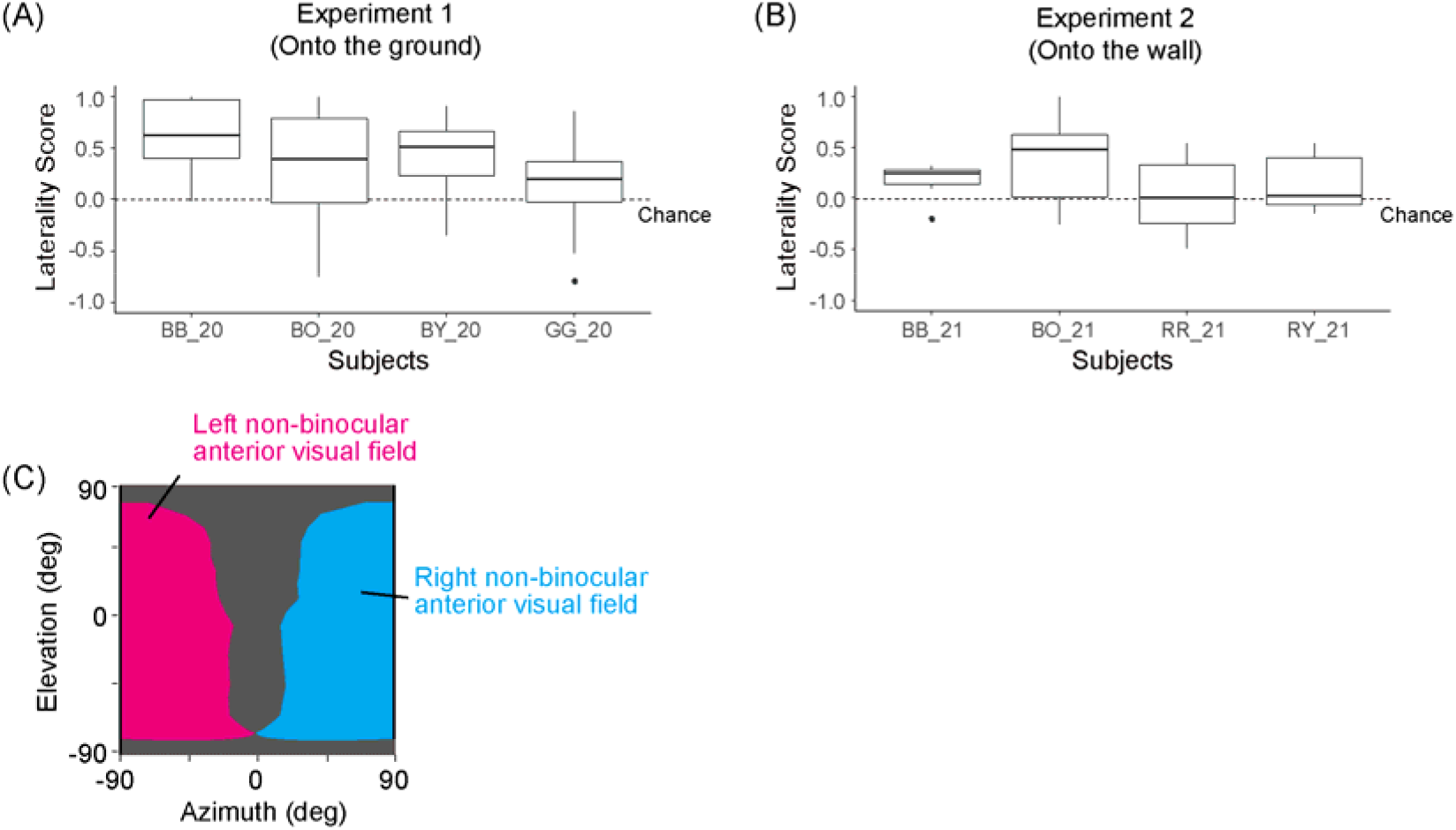
Laterality scores in each experiment. Laterality scores were defined as (right non-binocular anterior looking time-left non-binocular anterior looking time) / (right non-binocular anterior looking time + left non-binocular anterior looking time) in Experiments 1 (A) and 2 (B). Box plots show the median, interquartile range (IQR), and 1.5 × IQR, with outliers plotted individually. Only the sessions after visual targets landed on the ground or wall were tested because crows mainly use the non-binocular anterior visual field in Movement 3. The dotted line indicates the chance level (zero). The 2D visual field map created in Study 1 was used to calculate the laterality score (C). The right and left sides of the non-binocular anterior visual field are indicated in cyan and magenta, respectively.

## DISCUSSION

This study examined the visual field use of crows. Our main objectives were twofold: first, to identify the visual field regions they orient to when presented with an experimentally-defined attention-getting visual target, and second, to explore the contextual cues influencing the differential use of their left and right eyes, as well as distinct visual field regions. By addressing these objectives, we aimed to determine the feasibility of inferring a crow’s attentional focus from its head orientation. Study 1 examined the visual field configuration of large-billed crows, as it had not been identified in previous studies. We found that the binocular field covered approximately 50°, and eye movements ranged from approximately 16° to 17° in our crows when we did not induce eye movement. The optic axes were located at around 60° azimuth and −30° elevation.

With these visual field data, Study 2 employed a newly established motion capture system to investigate how freely moving crows orient their visual fields towards a distal (out-of-pecking) visual target. Crows used limited regions of the visual field, particularly the region around the beak projection (eye-beak-tip line), which is within the binocular region, as well as the regions around where we found the optic axes in Study 1, which is within the non-binocular region. The latter regions are likely where area *centralis*, the most visually sensitive spots in their retina, projects (Rahman et al., 2006).

In our experiments, the selective use of the binocular regions was mainly affected by the movement properties of the object. Specifically, when the experimenter wiggled the object, they tended to orient their binocular field to the object. When the experimenter threw the object, crows increased this tendency and typically followed the object’s movement via the smooth movement of their head (see Fig. 3). When the object landed on the ground or wall, crows tended to orient their non-binocular visual fields (likely their visual axes) to the object, typically via the saccadic movement of the head. They preferred to use the right eye when orienting their non-binocular visual fields to the object.

### Visual field configuration

How does the visual field configuration of this species compare to that of other corvid species? The maximum binocular overlap in this species was approximately 50°, regardless of whether the experimenter attempted to induce eye movements. This observation is largely comparable to those of other corvid species, which vary between 37.6° and 61.5° (Fernández-Juricic et al., 2010; Troscianko et al., 2012). The maximum eye movement amplitude was approximately 16–17° when we did not induce eye movement and 33° when eye movement was induced. The measurement of 16–17° (eyes at rest) is smaller than that of any other corvid species reported by Troscianko et al. (2012), who did not induce eye movement in corvids, but the measurement of 33° is largely comparable to that of other corvid species (22.2°–38.8°). Owing to large eye movements, the binocular field was temporarily lost when the eyes were attracted away from the ophthalmoscope, and a blind area was observed in the same visual field region. This observation is largely consistent with previous studies that tested several other passerine species, such as the American crow (Fernández-Juricic et al., 2010), American goldfinch (Baumhardt et al., 2014), chickadee (*Poecile carolinensis*), and titmouse (*Baeolophus bicolor*) (Moore et al., 2013).

### Visual field use

Why did crows prefer to use their binocular field when attending to an object in motion? Their binocular use is unlikely related to the use of area *centralis* because their optic axes (located at the azimuth of approx. 60°) cannot project into the binocular fields (50° in width, i.e., ±25° from the front), even when considering the largest eye movements observed in Study 1 (30°).

Rahman et al. (2006) reported that this species has a relatively high density of retinal ganglion cells in the dorso-temporal part of the retina, which projects forward. Thus, the binocular use of crows may be related to the relatively high quality of vision in their binocular field. An alternative possibility is that they use optic flow field information from the binocular field, such as the moving direction and time to contact a visual target. In birds, it is generally assumed that such information is obtained more efficiently from the binocular field than from the monocular field because the optic flow field appears symmetrical when a target object is located in the binocular field (Martin, 2009). Moreover, motion perception may be more fine-tuned in the binocular field because it may contain a relatively higher ratio of double to single cones in the retina, as found in several bird species (Fernández-Juricic et al., 2019; Tanaka, 2015).

### Right eye bias

Our crows were slightly biased toward using their right eye when viewing the presented visual target. Right eye bias (left hemispheric bias) is generally observed in birds when they attend to the detailed features of objects (Clayton & Krebs, 1994; Rogers, 2008). As our visual targets were small objects of various shapes and colors, our crows were likely motivated to attend to the detailed features of the objects. It should be noted that, in our experiments, the observed eye preference was weaker in Experiment 2 than in Experiment 1. One reason for this could be that the object tended to land on the ground in front of the crows in Experiment 1, while it landed on every side of the wall (approximately at the eye level of the crows); thus, it may have been physically easier for crows to select the preferred eye in Experiment 1. Thus, although our crows generally prefer to use their right eye, such a preference could easily be lost when there are certain physical constraints. In addition, although three out of the four crows showed a significant right eye bias in this study, it is important to note that previous research, including studies involving this species, has reported individual differences in the laterality bias among corvids (Clary et al., 2014; Izawa et al., 2005; Mack & Uomini, 2022). Therefore, caution is warranted when generalizing our results across individuals.

### Eye movement

We can assume that our motion capture system accurately tracked crows’ head orientations (Itahara & Kano, 2022). Thus, eye movements may largely explain variations in the distribution of objects in the visual field. Visual inspection of the heatmaps in Study 2, particularly Fig. 4A,B, suggests that eyes regularly shift the optic axes by approximately 15°, which is consistent with our observations in Study 1, where we observed 16–17° of eye movement when not attempting to induce eye movement (33° when inducing eye movement, possibly from one extreme angle to another). Fig. 4A,B indicates that the optic axis tends to shift both upward and downward in elevation but only inward in azimuth (the direction toward the beak). This suggests that crows may have mainly converged, rather than diverged, their eyes in our experiments. Typically, birds’ eyes converge when viewing a close object within their binocular field (Bloch et al., 1984; Troscianko et al., 2012).

Our additional analysis, shown in Fig. S4, examined the conditions under which crows might have used their convergent eyes to view a distant object with non-binocular visual fields. This analysis suggested that the peak in the heatmap was approximately 15° more inward when crows first followed the object with their binocular field and then oriented their visual axes (close to their optic axes) to the objects, compared to when crows viewed the object directly with their visual axes. This suggests that the crows first converged their eyes to view the object with the binocular field and then maintained these eye positions to fixate on the object near the optic axes. Overall, it appears that crows commonly exhibited eye movements within 15° in our experiments.

### Limitation of our tracking method

One limitation of our tracking method is that the motion capture system requires the attachment of a few lightweight reflective markers to the subject’s body. Although most of our subjects accepted the marker attachments, in Study 2, we could not test several subjects because they habitually removed the attached markers on their heads with their claws (see Methods). The use of a head-mounted camera (Kane et al., 2015; Kane & Zamani, 2014) or eye tracker (Schwarz et al., 2013; Yorzinski et al., 2015; Yorzinski & Platt, 2014) could significantly improve gaze-tracking accuracy, it may not be practical with corvid species, as they can easily remove such devices. Recent developments in marker-less tracking methods (e.g., Dunn et al., 2021; Nath et al., 2019; Pereira et al., 2022; Waldmann et al., 2022) offer potential solutions to the attachment issue. Nevertheless, it’s essential to carefully consider the trade-off between accuracy and applicability when choosing between infrared motion capture systems, which are generally more accurate, and marker-less tracking methods.

## Conclusion

Overall, our results indicate that crows exhibit predictable patterns for their visual field use, thus demonstrating the feasibility of inferring their attentional focus through 3D tracking of their head. Due to untracked eye movements, there is an estimated margin of error of 15° around their visual axis (Fig. 7). The differential use of their binocular and non-binocular fields likely depends on the motion of an object, according to our study, and also likely on the distance between the bird and an object, according to a previous study (Matsui & Izawa, 2019). The differential use of their left and right, although a weak tendency, may partly depend on the type of objects. Further investigation is required to explore what other contextual cues influence the differential use of their binocular or non-binocular fields and left and right eyes.

**Figure 7.**
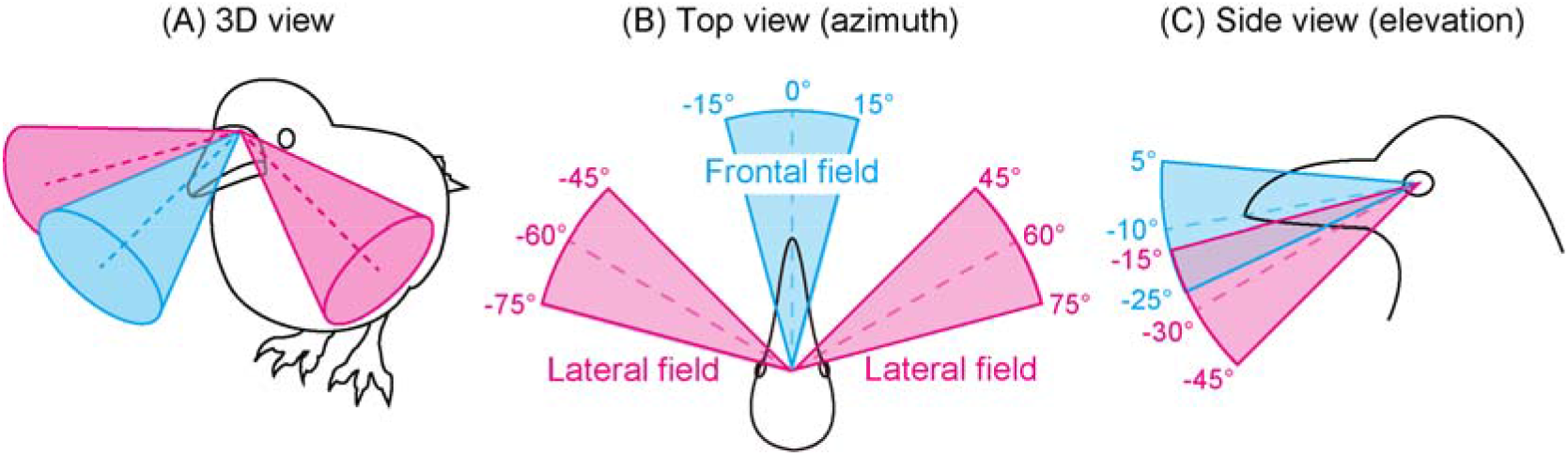
The simplified visual field model of large-billed crows. The frontal field around the projection of the beak-tip (eye-beak-tip line, cyan) and the lateral field around the optic axes (magenta) were drawn from oblique (A), top-down (B), and side views (C). Dotted lines indicate the angle of the beak-tip projection (in cyan) and optic axes (in magenta).

While more work is necessary, our current system represents a promising initial step towards establishing gaze tracking methods in the studies of corvid behavior and cognition. Notably, our specific procedures to test the possibility of gaze tracking can be applicable to other corvid and non-corvid bird species as well. Moreover, although this study tracked one bird at a time, our system can simultaneously track multiple individuals. As a result, it can be utilized to investigate socio-cognitive behaviors in birds, such as eye contact, gaze following, and theory of mind, within an interactive setup.

## ACKNOWLEDGMENTS

We thank the members of Kumamoto Sanctuary, especially Dr. Toshifumi Udono, for daily husbandry and veterinary care of the crows; Dr. Satoshi Hirata for the research support; Dr. Ei-Ichi Izawa for his advice on the visual field measurements; Dr. Hiroshi Matsui and Dr. Tomokazu Ushitani for the comments on the manuscript; Dr. James Brooks for English proofreading; the members of Ino-P Corporation for offering daily food for the crows; and Gyokuto-town for obtaining the crows.

## COMPETING INTERESTS

No competing interests declared.

## FUNDING

This work is supported by the Japan Society for Promotion of Science (KAKENHI 19H01772 and 20H05000), the Leading Graduate Program in Primatology and Wildlife Science, and the DFG Cluster of Excellence 2117 CASCB (ID: 422037984). We also thank Hayanon Science Manga Studio, Tomoichi Enomoto, and 78 people for supporting our research through the crowdfunding project on academist (https://academist-cf.com).

## DATA AVAILABILITY

All processed data, R codes for the statistics, the MATLAB codes for the data processing, and the sample data are available in OSF (https://osf.io/3dmg7/?view_only=c0f0a272ebff466ca8d0a3792da3ab4e).

The head calibration application is available on GitHub (https://github.com/itaharaakihiro/head_orientation_calibration_app_set).

## Notes

### Competing Interest Statement

The authors have declared no competing interest.

